# Unifying Single-Channel Permeability from Rare-Event Sampling and Steady-State Flux

**DOI:** 10.1101/2021.11.20.469392

**Authors:** Yi-Chun Lin, Yun Lyna Luo

## Abstract

Various all-atom molecular dynamics (MD) simulation methods have been developed to compute free energies and crossing rates of ions and small molecules through ion channels. However, a systemic comparison across different methods is scarce. Using a carbon nanotube as a model of small conductance ion channel, we computed the single-channel permeability for potassium ion using umbrella sampling, Markovian milestoning, and steady-state flux under applied voltage. We show that a slightly modified inhomogeneous solubility-diffusion equation yields a single-channel permeability consistent with the mean first passage time (MFPT)-based method. For milestoning, applying cylindrical and spherical bulk boundary conditions yield consistent MFPT if factoring in the effective bulk concentration. The sensitivity of the MFPT to the output frequency of collective variables is highlighted using the convergence and symmetricity of the inward and outward MFPT profiles. The consistent transport kinetic results from all three methods demonstrated the robustness of MD-based methods in computing ion channel permeation. The advantages and disadvantages of each technique are discussed, focusing on the future applications of milestoning in more complex systems.

## Introduction

Ion channels are complex biological nanopores that perform vital physiological functions with high sensitivity and precision. Over the decades, molecular dynamics (MD) simulation has become an indispensable tool for computing the functional properties of ion channels directly from their dynamic structures. Various MD-based methods were developed for investigating the thermodynamics and kinetics of ions or small-molecules permeation at the single-channel level. Many pioneering atomistic MD simulations on ion channels have focused on computing ion permeation from equilibrium free energy profiles, or potential of mean force (PMF), in conjunction with electro-diffusion theory^1-4^. The increased computing power and performance of MD engines have also enabled researchers to simulate single-channel conduction explicitly under a constant external electric field^5, 6^ or an asymmetric ionic concentration across the channel^7, 8^. If the system reaches a steady-state under voltage or concentration gradient, a mean flux rate and a steady-state density profile can be obtained from the ensemble of nonequilibrium processes. These equilibrium and nonequilibrium MD simulations have significantly deepened our understanding of the ion channel permeation process at the high temporal and spatial resolution, as reviewed by ref^9-12^.

Unlike the steady-state flux under voltage, the equilibrium MD approaches can be generally applied to any small-molecule permeation (neutral or charged). Several theoretical frameworks can be used to compute crossing rates from PMF profiles obtained from enhanced sampling simulations. Particularly, if the PMF is dominated by a single large barrier and the permeant diffusion is constant at the barrier region, crossing rate can be estimated via Kramer’s theory or transition state theory (TST) borrowed from reaction kinetics. However, for complex biological ion channels, the aforementioned assumptions may be far from satisfied. Alternatively, molecule permeation may be considered as a one-dimensional nonreactive diffusive process that can be described using the fluctuation-dissipation theorem. For instance, PMF can be used together with position-dependent diffusion coefficient to estimate permeability using inhomogeneous solubility-diffusion (ISD) equation^13^. ISD equation has been applied successfully in studying solute permeation across membrane^14, 15^. Here we show that a slightly modified ISD equation yields a single-channel permeability that is consistent with a mean first passage time (MFPT)-based method, which extracts detailed kinetics along the molecular permeation pathway directly from rare-event sampling trajectories. Recent examples of such rare-event sampling applied on ion channels include Milestoning^16-18^, Weighted ensemble sampling^19^, and Markov state models^20-23^. In theory, the PMF-based method, MFPT-based method, and steady-state flux under voltage should converge to the same single-channel permeability for the same studied system. However, a systemic comparison between different methods is still lacking.

In this work, we use a carbon nanotube (CNT) as a model (**Figure 1a**) of small conductance (∼2 pS) ion channel to compare K^+^ permeability from milestoning, umbrella sampling (US), and voltage simulations. This CNT system has been used to compute K^+^ permeability using a transition path approach similar to the reactive flux method^24^. We chose this system because its free energy barrier height (∼4 kcal/mol) and microsecond-timescale crossing rate are physiologically relevant. Such a system requires nontrivial sampling (beyond the capability of brute-force MD), but the rigidity of the CNT still allows good convergence and unambiguous comparison of all methods tested here.

**Figure 1.**
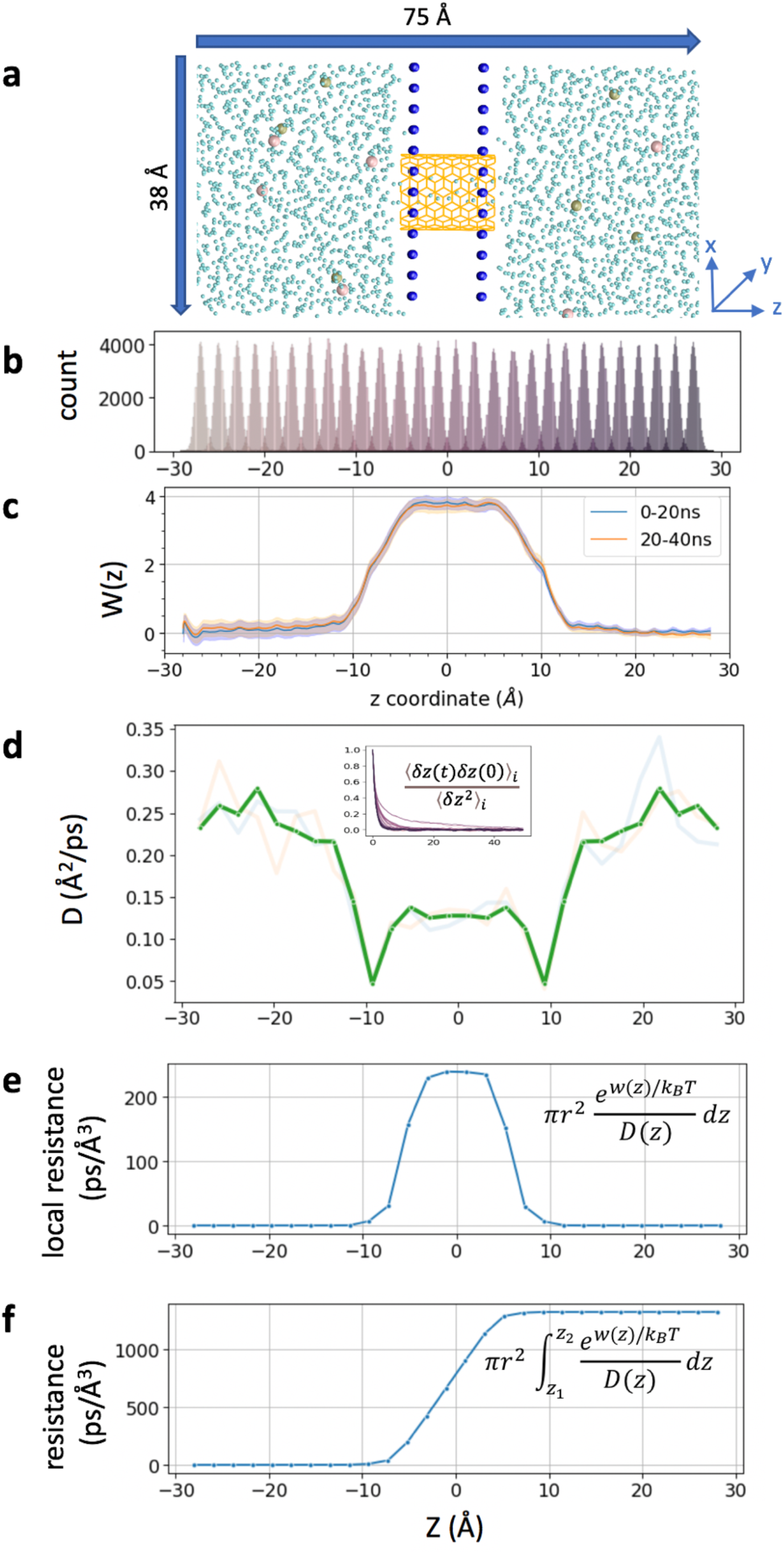
**a**. Simulated CNT system, consisting of two fixed layers of carbon atoms as water impermeable membrane. 6 K^+^ and 6 Cl^-^ ions are shown in tan and pink, CNT in orange color, and water oxygen in cyan. Membrane carbon atoms are shown as blue spheres. **b**. z-coordinate distribution of a tagged K^+^ from each umbrella sampling window (corresponding data from milestoning is shown in Figure 2b). **c**. PMFs from two blocks of umbrella sampling trajectories, generated from MBAR with 80,000 data points per window, with error bars in shaded color. **d**. The position-dependent diffusion constant of K^+^ computed from umbrella sampling. The transparent lines represent data from the first 20ns and second 20ns per window. A thick green line represents averaged and symmetrized values. Inset is the plot of correlation function used to compute correlation time *i* and *D(z)* (see methods). **e**. Local resistance (the integrand of Eq 3) for permeating K^+^. **f**. Integration of the permeation resistance, 1/*P*, as a function of z.

The original milestoning simulation requires running short trajectories in each milestone until they reach another milestone^**25**^. Here, we make use of the “soft-walls” Voronoi-tessellated Markovian milestoning, which confines the sampling within the Voronoi cells using flat-bottom harmonic restraining potentials^**26**^. The implementation of this “soft-walls” version (referred to as milestoning thereafter) resembles, to a large degree, the conventional umbrella sampling setup. A detailed comparison of the sampling, PMF, and MFPT results from milestoning and umbrella sampling is the focus of this study. In addition, we tested two bulk boundary conditions, namely the cylindrical and spherical boundaries, that are particularly useful for conducting milestoning on ion channels.

The CNT system chosen here is designed to satisfy the symmetric single barrier requirement, thus allowing us to check the robustness of the milestoning method by computing both inward and outward permeation rates. We also show that the MFPT from milestoning is extremely sensitive to the frequency of recording the relevant collective variables (e.g., the coordinates of the tagged ion). All physical quantities and the obtained results are summarized in **Table 1**. The overall consistent single-channel permeability demonstrated the robustness of the theoretical and computational framework tested here. The limitation and strengths of each method are discussed and compared.

**Table 1.** Summary of the three methods for computing single-channel permeability.

| Method | Umbrella sampling | Markovian milestoning | Steady-state flux |
| --- | --- | --- | --- |
| Permeability equation | $P = \pi r^2 \left( \int_{z_1}^{z_2} \frac{e^{w(z)/k_B T}}{D(z)} dz \right)^{-1}$ | $P = \frac{\pi r^2 \int_{z_1}^{z_2} e^{-w(z)/k_B T} dz}{2 \times MFPT}$ | $P = \frac{\gamma k_B T}{q^2 C}$ |
| Effective bulk conc. | 0.77M | 0.77M | 0.52 M |
| Biasing potential | Harmonic | Flat-bottom harmonic | Voltage $\pm 0.4 \text{ V}$ |
| Biasing force constant | 2.5 kcal/mol | 100 kcal/mol | - |
| Lateral restraint in bulk | 6 Å | 6 Å | - |
| PMF barrier (kcal/mol) | 3.8±0.17 | 4.1±1.7 | - |
| Key parameters | D(z) 0.004 ~0.028 Å<sup>2</sup>/ps | MFPT: 2.6 μs | conductance 2.3±1.2 pS |
| Permeability (cm<sup>3</sup>/s) | (8.96±0.02) $\times 10^{-16}$ | 2.92 $\times 10^{-16}$ | (11.7±6.36) $\times 10^{-16}$ |
| Total sampling time (μs) | 1.12 | 4.98 | 0.60 |

## Theory and Methods

### Relation between single-channel permeability, MFPT, and conductance

Assuming permeating molecules do not interact under sufficient low concentration, under physiological conditions, single-channel permeability *P*(cm^3^/s) can be related linearly to the rate of crossing *k* or mean first passage time (MFPT) ⟨*t*⟩under equilibrium, *P*= *k*/*c*= 1/*c*⟨*t*⟩, in which *c*is the symmetric solute concentrations. Here, we use the number of molecules per second for *k*, seconds per molecule for MFPT, and molecule/cm^3^ for *c*.

For ionic permeation under voltage and/or concentration gradient, Goldman-Hodgkin-Katz (GHK) flux equation describes the ionic flux across a homogenous membrane as a function of a constant electric field (voltage) and an ionic concentrations gradient. Under symmetric concentration and constant voltage, the current (*I*) and the permeability (*P*) can be related by the GHK flux equation, 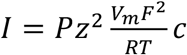, where *z* is the charge of the permeant, *V*_*m*_ is the voltage, *F* is the Faraday constant, *R* is the gas constant, *T* is the absolute temperature, and *c*is the concentration^27^. When applied to a membrane-embedded single-channel model and ions only cross the membrane through the channel, *P*in the GHK equation corresponds to the single-channel permeability. GHK flux equation thus relates single-channel conductance *γ*, a nonequilibrium property, to the equilibrium property *P*.

It can be seen from the equations above that the crossing rate *k*, MFPT, and conductance *γ* are all concentration-dependent; only *P* is independent of concentration. Experimentally, *P* is usually measured relative to the potassium ion permeability, thus representing an intrinsic property of each channel. It is hence an ideal quantity for rigorous comparison between different computational methods.

### System setup and equilibrium protocols

The coordinates of the carbon tube (CNT) were taken from ref^24^. Briefly, it is an uncapped armchair CNT with 13.5 Å in length and 5.4 Å in radius. Two carbon sheets form an artificial membrane to separate the solution (**Figure 1a**). Constraints were applied to all carbon atoms to keep the system rigid. CHARMM36 force field was used^28, 29^. After solvation, the box size was 38 ×38×75 Å^3^, which contained 2503 TIP3P water molecules, six K^+^ and six Cl^-^. All MD simulations were performed using 1 fs time step using NAMD2.13 package under NVT ensemble, with 1 atm and 300 K temperature using Langevin thermostat^30, 31^. Cutoff for calculating vdW interaction and short-range electrostatic interaction was set at 12 Å and force-switched at 10 Å. Long-range electrostatic interactions were calculated using the particle mesh Ewald algorithm^32^. The system was equilibrium for 100ns before conducting umbrella sampling, milestoning and voltage simulations.

### Umbrella sampling simulations

A total of 28 windows (−27 Å < z < +27 Å) were sampled. Each window was separated by 2 Å apart. The tagged K^+^ was restraint by a harmonic restraint on z and a flat-bottom harmonic cylindrical restraint. The force constant for harmonic restraint was 2.5 kcal/mol/Å and cylindrical restraint was 10 kcal/mol/ Å within 6 Å on X-Y radius plane. The reference dummy atom to pull the K^+^ was set at (0,0,0). The harmonic distance restraint was determined by the projected vector along z between the dummy atom and tagged K^+^. The cylinder restraint was determined by the center of mass of all carbon atoms from CNT. Each window was run for 40 ns (**Figure 1b**). The PMF (**Figure 1c**) was computed using pymbar 3.0.3^33, 34^. The output frequency was 0.5ps per frame.

### Position-dependent diffusion coefficient D(z)

D(z) of K^+^ inside the CNT was calculated from umbrella sampling trajectories (**Figure 1d**). The correlation time was extracted from each umbrella window *i* using 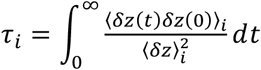, where *δz*(*t*) = *z*(*t*) −⟨*z*⟩, is the deviation of the *z*-position of the ion at time *t, z*(*t*), from the time-averaged position ⟨*z*⟩ in each window.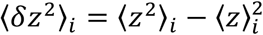, is the variance. Following the formulation of ref^35-37^, the Laplace transformation of the velocity autocorrelation function along the reaction coordinate *z* in the harmonically restrained umbrella sampling gives 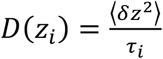.

### Markovian Milestoning with cylindrical bulk restraint

Same as in umbrella sampling, the z-coordinate of the tagged ion was used to define a set of Voronoi cells along the channel pore, and to identify the milestones as the boundaries between the cells. To facilitate the comparison in analysis, we kept the Voronoi cell setup (28 cells and 2 Å apart) and the bulk cylindrical restraint (6 Å radius) identical as our umbrella sampling windows (**Figure 2 a**). The only difference is that a flat-bottom harmonic restraint of force constant 100 kcal/mol/Å, instead of the weak harmonic restraint, was used to confine the sampling within each cell (**Figure 2b vs. Figure 1b**). We then ran 28 local simulations confined in each cell, and collected the kinetics of transitions between milestones. More specifically, let us introduce a set of *M* Voronoi cells *B*_i_, *i*=1,*…,M*. Since the total flux in and out of each cell is zero at statistical equilibrium, the rate of attempted escape from cells *B*_*j*_ to *B*_i_, *k*_*j* →*i*_, and the equilibrium probability π_i_ for the tagged ion to be in cell *B*_i_ satisfies a balance equation:

**Figure 2.**
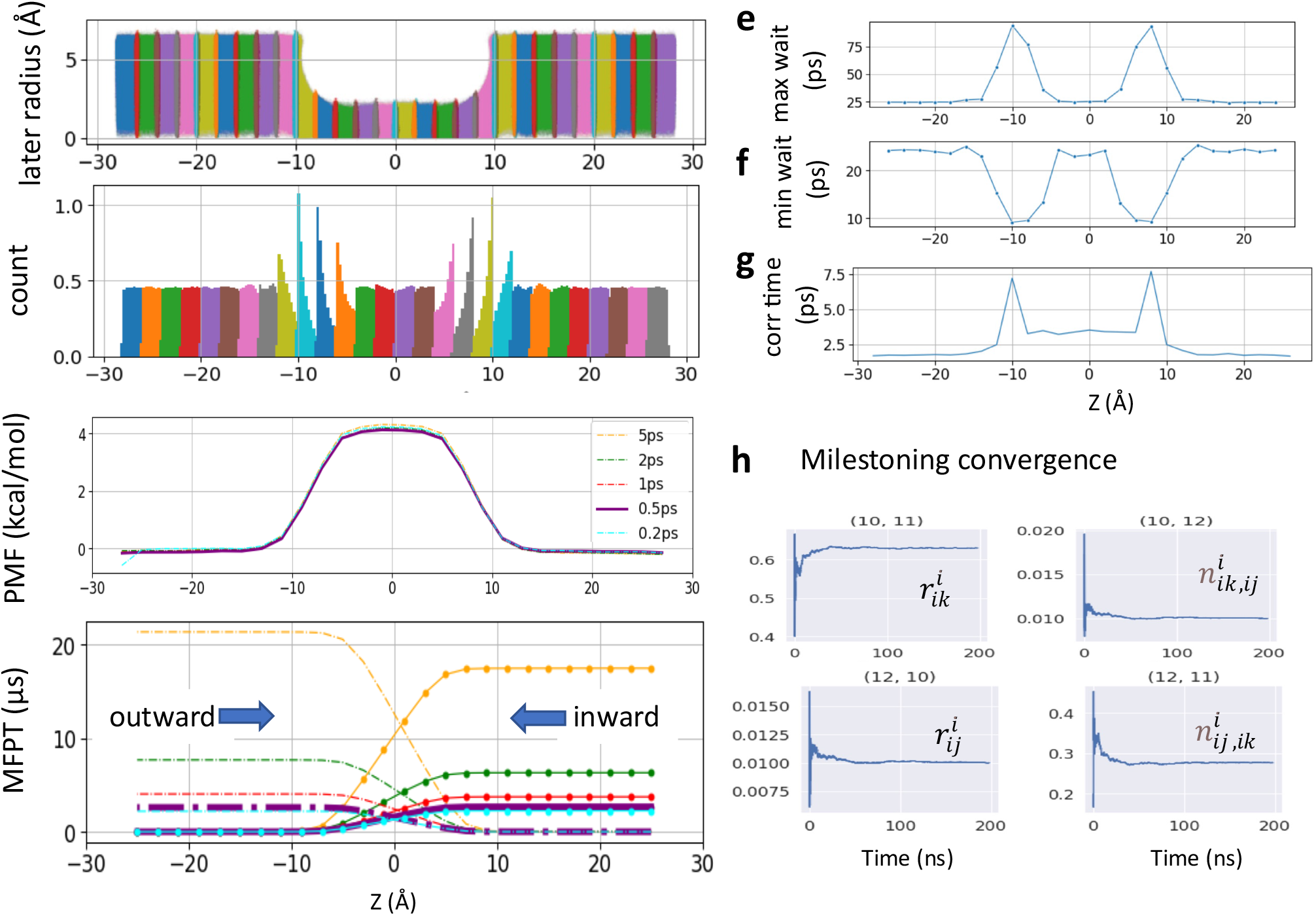
**a**. Raw data plotted along the channel z-axis and radial distance 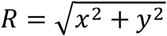 from channel center axis (*x,y* = 0,0) from 28 milestoning sampling cells. **b**. Distribution of the milestoning data along the z-axis (same plot from the US is shown in Fig1a). **c**. PMF from Milestoning sampling. Different color represents different Colvars frequency (i.e., the frequency of recording the z-coordinates of the tagged ion). The bold purple (0.5 ps) is the data used for the final comparison. **d**. MFPT plot with the same color representation as PMF plot. The curves with dot solid lines are outward directions of tagged ion, and dash curves present inward direction. **e**. Maximum waiting time between successive transitions. **f**. Minimum waiting time between successive transitions. **g**. z-position decorrelation time in each milestoning cell. **h**. Convergence of the variables in Eq. 2 for computing the rate matrix. Milestoning cell 11 is chosen as an example here.

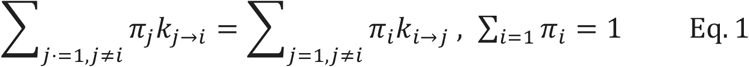

The free energy of each cell can be obtained from the solution of **Eq. 1** as −*k*_*B*_*Tln*(*π*_i_) (**Figure 2c**). By defining a milestone *S*_ij_ as the boundary between two adjacent Voronoi cells *B*_i_ and *B*_j_, the dynamics of the system is reduced to that of a Markov chain in the state space of the milestone indices^38^. The MFPT between any pair of milestones *S*_ij_ and *S*_ik_ can hence be calculated from the rate matrix whose elements *q*_*ij,ik*_, the rate of moving from milestone *S*_ij_ to *S*_ik_, are given by **Eq. 2**:

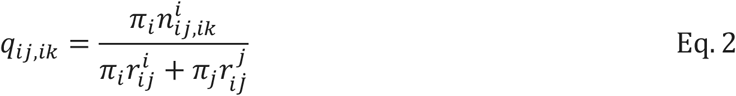

where 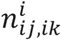 is the number of transitions from *S*_ij_ to *S*_ik_, normalized by the time spent in cell *B*_i_. 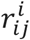 is the time passed in cell *B*_i_ after having hit *S*_ij_ before hitting any other milestone, normalized by the total time spent in cell *B*_i_. The inward and outward MFPT profiles were obtained by reversing the milestone indices when constructing the rate matrix (**Figure 2d**). The 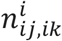 and 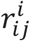 can be used to monitor the convergence of the rate matrix (**Figure 2h**).

The total sampling time of all 28 cells was 4.9 µs, where each cell was sampled between 150 ns to 300 ns. The NAMD Colvars output frequency was 0.5ps. The PMF and MFPT were computed using a set of in-house python scripts https://github.com/yichunlin79/CNT_milestoning_method with different frame sizes. To check whether the Colvars output frequency has any effect on MFPT, additional milestoning simulations were conducted with the Colvars output frequency of 0.2 ps and a total sampling time of 2.74 µs.

### Markovian Milestoning with spherical bulk restraint

For spherical bulk restrained milestoning, a total of 14 Voronoi cells were used, including eight cells inside the channel (identical to the milestoning above) and three layers of spherical shell on each side of the channel (**Figure 3a)**. The distance between the tagged K^+^ ion and two dummy atoms fixed at the cartesian coordinates of (0,0,-8) and (0,0,8) are used to set up the spherical shells in bulk with radius increments of 3, 3, and 4 Å. Additional z>|8|Å restraint is applied to keep the ion outside the channel. All restraint force constant is 100 kcal/mol/Å. The length of each bulk window is 150 ns with Colvars output frequency of 0.5 ps^-1^.

**Figure 3.**
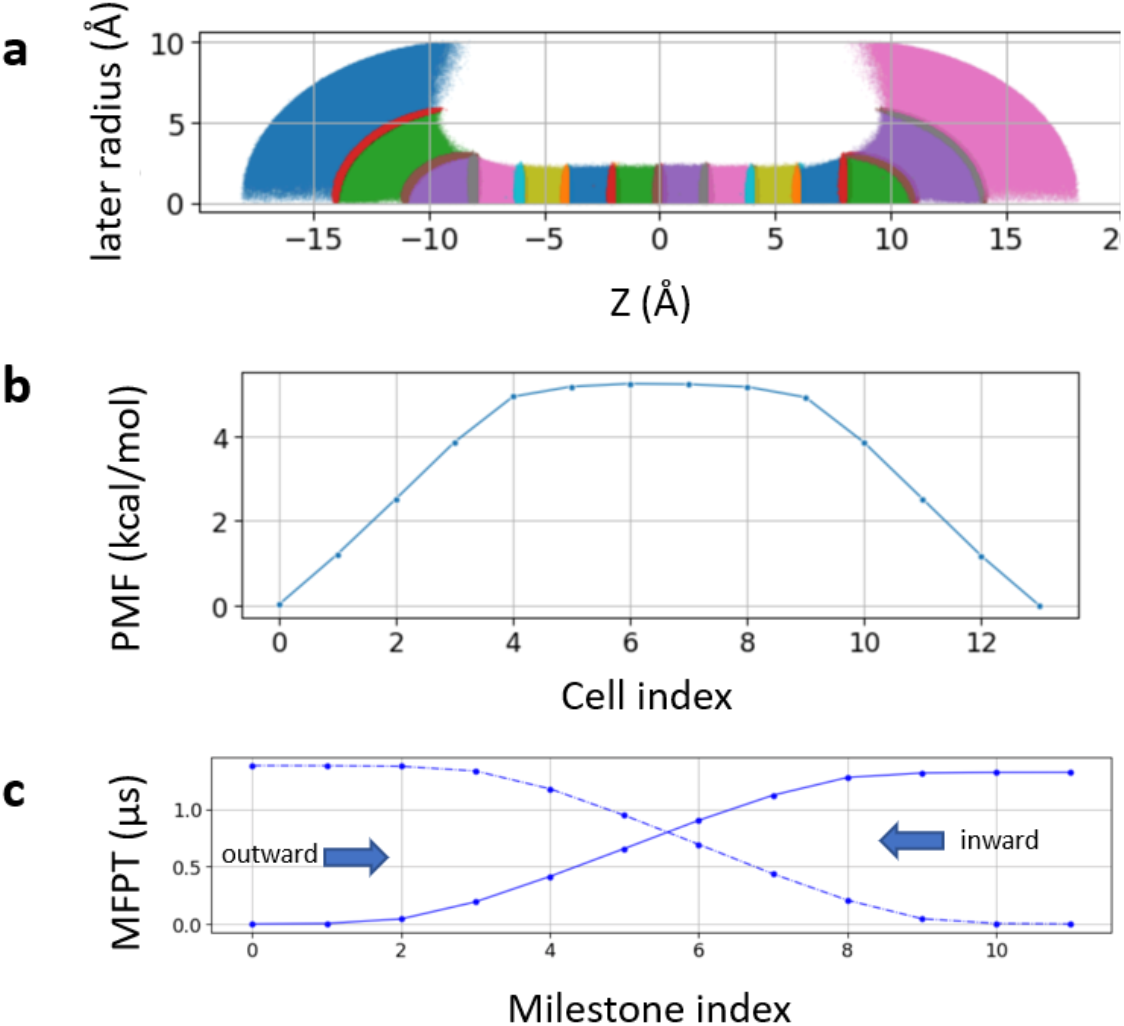
**a)**. Sampling plot with spherical restraint at two bulk ends. The three spherical radius intervals are 3 Å 3 Å and 4 Å from channel to bulk. **b)**. PMF of the spherical restraint system. The x-axis represents the cell index number. **c)** MFPT of the spherical restraint system. X-axis represents the cell index numbers.

### Voltage simulations

After 100 ns equilibrium simulation, constant electric fields corresponding to the transmembrane potential of +/-0.3 and +/-0.4 V were applied perpendicular to the membrane to all the atoms using NAMD2.13. To be consistent with the umbrella sampling and milestoning, a cylinder restraint of 6 Å radius was applied to a tagged K^+^ in bulk with 10 kcal/mol/ Å force constant for +/-0.4V systems. All other ions moved freely beyond the cylinder restraint region. The K^+^ conductance was computed by counting the total number of crossing events and by computing the charge displacement along the z-axis (**Figure 4**). Error bar was computed from three independent replicas of 200 ns. For +/-0.3V systems, all six K^+^ were restrained inside the same cylindrical bulk boundary and a single replica of 60 ns was conducted. The time step was 1 fs and the output frequency was 0.5 ps for all voltage systems.

**Figure 4.**
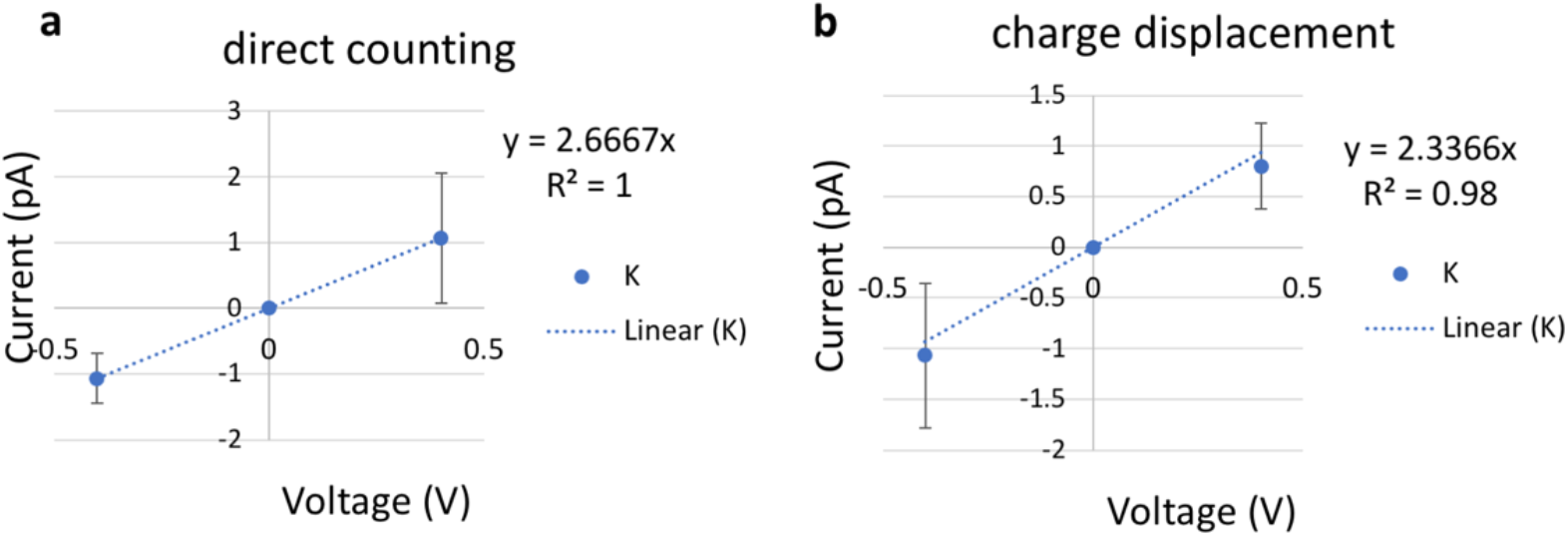
current-voltage (I-V) plot for a single K ^+^ with a cylinder restraint of radius 6 Å in bulk. The slope of the linear fitting defines the conductance. **a**. K^+^ current computed using charge displacement. **b**. K^+^ current computed using direct counting method. The error bars are calculated from three replicas of 200 ns at each voltage.

## Results and Discussion

### Single-channel permeability from inhomogeneous solubility-diffusion equation (ISD)

Molecular permeation through ion channel can be described by ISD if the one-dimensional free energy and diffusion along channel normal is sufficient to describe the diffusion process (relaxation of orthogonal degrees of freedom is fast relative to the reaction coordinate) and the permeant velocity relaxation time is instantaneous (on the scale of integration time step). The only difference with the ISD equation used for membrane permeation is that a flat-bottom lateral potential *u*(*x, y*) is often used to confine a single tagged ion in a cylindrical bulk region outside the ion channel. The effective cross-sectional area due to the lateral restraint is thus ∬ *e*^− *βu*(*x,y*)^*dxdy*, which can be approximated to *πr*^2^ in a homogenous bulk, where *r* is the radius of the cylinder. Hence, *πr*^2^ defines the effective bulk concentration in the simulated region, which led to the probability of the ion inside the channel *p*(*z*) over the true bulk density *ρ* to be 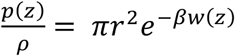, where *w*(*z*) is the PMF with the bulk value set to zero at the cylindrical region and *β* = 1/*k*_*B*_*T* ^39, 40^. *T* is the temperature; *k*_*B*_ is the Boltzmann’s constant. Therefore, at low ionic concentration, single-channel permeability can be estimated using a slightly modified ISD equation,

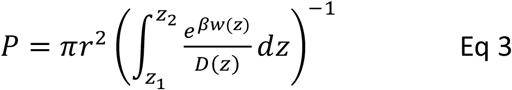

In **Eq. 3**, *D*(*z*) is the position-dependent diffusion constant of the studied permeant along the *z*-axis (**Figure 1d**). The interval of the integration, [*z*_1_, *z*_2_], are the lower and upper boundaries of the channel pore, beyond which PMF reaches the bulk value. *r* is the radius of the cylindrical restraint when ion is outside of [*z*_1_, *z*_2_] interval. It is necessary to set *r* to be larger than the maximum pore radius so that it has no energetic contribution inside the pore. The radius of the cylindrical restraint defines the effective bulk concentration. Thus, it offsets the bulk PMF value and ensures that the single-channel permeability from **Eq. 3** is concentration-independent.

In both umbrella sampling and milestoning, same cylindrical restraint with *r* =6 Å is applied in the bulk region, and the same window size of 2 Å was used. The only difference is that a weak harmonic restraint with force constant of 2.5 kcal/mol is applied for all umbrella windows to ensure sufficient overlapping between neighboring samplings, but a strong flat-bottom harmonic restraint with a force constant of 100 kcal/mol is applied for all milestoning cells to confine the sampling within each cell. **Figure 1b** and **Figure 2b** illustrate the biased sampling distribution imposed by these two types of restraints. The PMFs from the US are shown in **Figure 1c**. With bulk value offset to zero, a broad energy barrier of 3.8 kcal/mol located inside the channel region is consistent with a previously reported PMF^24^.

Using the PMF or *w*(*z*) in **Figure 1c** and D(*z*) in **Figure 1d**, the permeability estimated from **Eq.3** is (8.96± 0.02) × 10^−16^ cm^3^ /s. We can also plot local resistance (the integrand of **Eq. 3**) for permeating K^+^ through CNT (**Figure 1e**) and the integration of the permeation resistance, 1/*P*, as a function of the z-axis (**Figure 1f**). It is not surprising that the 1/*P*(*z*) bears the same feature as the outward MFPT in **Figure 2d**.

### Permeability computed from mean first passage time (MFPT)

The MFPT of a single K^+^ crossing the CNT is computed from Voronoi-tessellated Markovian milestoning simulations (see Methods). The distributions of the tagged K^+^ confined in each 2 Å cell by flat-bottom harmonic restraint are shown in **Figures 2a** and **2b**. Milestoning simulation yields a consistent PMF profile with the highest energy barrier of 4.1 kcal/mol at the center of the CNT (**Figure 2c**). Since the CNT used here is symmetric by design, a rigorous check of sampling convergence is the perfect symmetricity (mirror image) of the inward and outward MFPT profiles (**Figure 2d**). The inward and outward MFPT profiles can be obtained by reversing the milestone indices when constructing the transition rate matrix.

We found that while PMF is nearly insensitive to the Colvars frequency (i.e., the frequency of recording the z-coordinates of the tagged ion) tested here, MFPT is susceptible to this frequency. In the current study, the frequency of 0.5 ps^-1^ severely overestimates the MPFT due to the missing transition events. Lower frequency also yields less data, which leads to asymmetric MFPTs. Here, the MFPT from the sampling saved per 5 ps has ten times fewer data points than the one from 0.5 ps, thus it failed to converge even after 18 µs of sampling. The ideal frequency has to be system-dependent (local diffusion and shape of the underlying free energy landscape). For our CNT system, the Colvars frequencies of 0.2 ps^-1^ and 0.5 ps^-1^ yield an identical and symmetric MFPT of 2.6 ± 0.03 µs for K^+^ permeation.

Using the PMF, *w*(*z*), and MFPT, ⟨*t*⟩, from the milestoning (with 0.5 ps^-1^ Colvars frequency), the single-channel permeability computed from **Eq. 4**^41^ is 2.92 × 10^−16^ cm^3^ s^-1^, in fairly good agreement with the permeability of 8.96 × 10^−16^ cm^3^s^-1^ computed from ISD equation (**Eq. 3**) using umbrella sampling data.

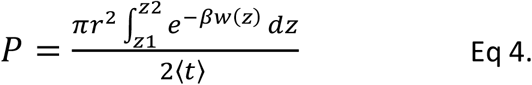

### Decorrelation time vs. waiting time in milestoning

Vanden-Eijnden et al. have shown that Markovian Milestoning yields exact MFPTs if the milestones are chosen such that successive transitions between them are statistically independent^38, 42^ and thus, no definition of lag time is needed. To check this assumption for transitions between two neighboring milestones, in each cell, the maximum and minimum waiting time between two neighboring milestones is extracted and plotted in **Figures 2e** and **2f**. The mirror image-like relation between these two plots manifests the fact that the transition down the slope of PMF is the fastest, and the one against the slope is the slowest. Hence, the longest waiting time 94.0 ps and the shortest waiting time 9.0 ps are in the same cell where the PMF is steepest. The velocity decorrelation time is less than the smallest frame size (0.2 ps). The z-position decorrelation time for the tagged K^+^ in each cell is plotted in **Figure 2g**. The maximum positional decorrelation time (7.6 ps) is also located at the steepest PMF region. Hence, for all pairs of milestones, the velocity decorrelation time and position decorrelation time are both less than the minimum waiting time for successive transitions between milestones.

### MFPT computed from spherical boundary condition

Laterally confining the ion in the bulk region (cylindrical restraint) is convenient for describing the thermodynamics and kinetics of the ions across the channel along the channel pore axis (z-axis). However, the geometries of ion channels are diverse. For funnel-shaped channel pores (e.g., connexin hemichannel^43^) or pores connected with lateral fenestration (e.g., Piezo1 channel^44^), a spherical boundary may be a better choice to capture the distribution and dynamics of ions near the channel entrance. Hence, we further tested milestoning simulation using spherical restraint for ions in the bulk region (**Figure 3a**). Unlike the cylindrical restraint, which yields constant ionic concentration along the z-axis, the effective ionic concentration in the current spherical bulk cells decreases as the radius of the sphere increase. Thus, the PMF and MFPT are plotted against the milestoning cell index rather than the z-axis (**Figure 3bc**).

At low concentration, single-channel permeability (cm^3^ /s) can be related linearly to the mean first passage time (MFPT) ⟨*t*⟩ under equilibrium P=1/c⟨*t*⟩, in which *c* is the symmetric solute concentrations. Since single-channel permeability is an intrinsic property of a channel, independent of solute concentration or the shape of the bulk cells, the ratio of MFPT from spherical restraint over cylindrical restraint should be equal to the reciprocal bulk concentration ratio. The concentration of a single ion in a hemisphere with a radius of 10 Å is 1.26 M and in a cylinder with a radius of 6 Å and length of 10 Å is 0.68 M. Therefore, the concentration ratio of ∼2 is indeed consistent with the MFPT of 2.6 µs from cylindrical restraint (**Figure 2e**) and 1.3 µs from spherical restraint (**Figure 3c**).

### MFPT computed from umbrella sampling

In the high diffusion limit, the MFPT of diffusive motion of K^+^ from the lower to upper boundaries of the channel pore, [*z*_1_, *z*_2_], can be written as

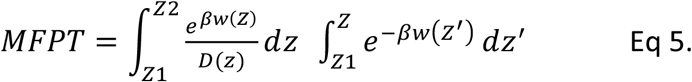

**Eq 5** was originally developed for computing the average reaction time for diffusion processes governed by a Smoluchowski-type diffusion equation^45^. It is also used to derive **Eq 4** from **Eq 3**^41^. To cross-validate our results, we apply the *w*(*z*) and *D(z)* from umbrella sampling to **Eq.5**, and obtain an MFPT of 2.2±0.02 *µ*s, fairly similar to the 2.6 *µ*s MFPT computed directly from milestoning. This consistency further demonstrated the robustness of the tested MD methods for computing transport kinetics.

### Permeability computed from steady-state flux

As mentioned above, under low concentration and constant electric field, we can simplify the GHK flux equation for computing the single-channel permeability *P*from conductance measurement. Under symmetric concentration, the GHK flux equation can be written as **Eq.6**.

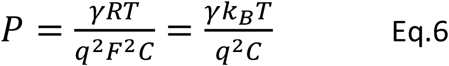

where *γ* is the unitary conductance of a single channel, *R* is the gas constant, *q* is the charge of the permeating ion, *F* is the Faraday constant, *C* is the bulk concentration of the ion, *k*_*B*_*T* has the same meaning above, except the unit is eV here (0.026 eV at 300 K). At sufficiently small voltage, the current-voltage (I-V) relation is expected to be linear, and the slope defines the conductance.

Ionic conductance from MD simulations can be computed using two approaches. The most commonly used is a direct counting method, in which the currents were computed from the number of permeation events (*N*) over a simulation period (*τ*), *I=N*/*τ*. In our code (see Github link in Methods), the channel is split into upper, inner, lower regions. A positive permeation event of K^+^ is counted if the time evolution of ion coordinates follows a lower-inner-upper sequence, and a negative permeation event is in reverse order. The carbon nanotube is applied with positive and negative 0.4 V voltage with 200ns per replica. Each voltage simulation was repeated three times with different initial velocities. A least-square fitting of the I/V curve gave the conductance of 2.7±0.94 pS by the direct counting method (**Figure 4 a**).

A more efficient approach that does not rely on completed permeation events is to compute the instantaneous ionic current from charge displacement along the z-axis, 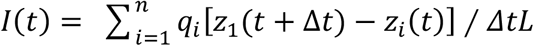, in which *q*_*i*_ and *z*_*i*_ are the charge and z coordinate of ion *i*, and *L* is the length of the channel pore^46^. This charge displacement method yielded a similar conductance of 2.3±1.2 pS, indicating a good convergence of the voltage simulations (**Figure 4b**).

With an ionic concentration of 0.52 M (one single K^+^ in a cylinder bulk of radius 6 Å and length 28 Å), the permeability is (11.9±6.36) × 10^−16^ cm^3^/s by charge displacement method, and (13.6±4.81) ×10^−16^ cm^3^/s by direct counting method. In addition to +/-0.4V simulations, ionic current calculated from 60 ns of +/-0.3 V with 6-fold higher concentration yields a similar permeability of 10.9 × 10^−16^ cm^3^/s. Only the results from low concentration are reported in the summary **Table 1**.

## Discussions

In this study, we used a carbon nanotube (CNT) as a toy model of a small conductance ion channel and computed the single-channel permeability from umbrella sampling, Markovian milestoning, and steady-state flux under voltage (**Table 1**). The PMF and MFPT for a single K^+^ permeating through the CNT produced from Markovian milestoning and umbrella sampling are in good agreement. Milestoning with cylindrical bulk restraint and spherical bulk restraint were tested and yielded consistent MFPTs when the effective bulk concentration is accounted. The single-channel permeability from voltage simulation is also within the same order-of-magnitude of those obtained from PMF-based and MFPT-based methods. These results are also in the same range as the previously reported K^+^ permeability of (25±7) ×10^−16^ cm^3^ /s computed using a transition path approach with CHARMM22 force fied^24^.

One caveat of the steady-state flux simulation under voltage is that the conductance estimated from the slope of the I/V curve assumes a linear response of the channel crossing rate. Even though +/- 400 mV is routinely used for MD simulations of ion channels to accelerate ion crossing, it may overestimate conductance due to the non-linearity of the I/V curve beyond physiological voltage. The non-linearity can be more prominent in a small system like CNT. Thus, the single-channel permeability reported from the steady-state flux simulation should be considered as an upper limit when compared with the permeability obtained from equilibrium approaches.

It should also be noted that the current CNT model is chosen because it reproduces macroscopic properties (e.g., free energy, conductance) similar to common small-conductance ion channels. However, two carbon sheets were used as an artificial membrane to separate the solution. In the absence of a dielectric medium surrounding the channel transmembrane region, this toy model is hence not suitable for investigating detailed electrostatic interaction between the ions and the channel.

In terms of computational resources, umbrella sampling has the advantage because PMF converges much faster than MFPT. However, MFPT allows extracting the kinetics directly from sampling, thus does not rely on the assumption of ISD formulism and does not require additional calculations of position-dependent diffusion coefficient. Steady-state flux is straightforward to apply if the permeant is charged and sufficient sampling is achievable under reasonable voltages. However, if the permeant is not charged, a constant concentration gradient needs to be applied. Furthermore, the effect of an unphysiologically large voltage bias or electrochemical gradient on channel property is likely system-dependent and difficult to predict over a long simulation time. Compared with the steady-state flux approaches, the milestoning approach does not depend on the charge of the permeant. Thus, it can be used to study any type of small molecular permeation, such as the transport of a second messenger, cAMP, through a connexin26 hemichannel^43^. Therefore, the use of milestoning has a significant promise for future applications on complex systems that are difficult to extract kinetics from unbiased MD or PMF-based enhanced sampling approaches.

## Acknowledgment

We are grateful to Dr. Fangqiang Zhu for sharing the coordinates of the CNT model, Drs. Luca Maragliano and Andrew Harris for inspiring discussions. This work was supported by NIH Grants GM130834. Computational resources were provided via the Extreme Science and Engineering Discovery Environment (XSEDE) allocation TG-MCB160119, which is supported by NSF grant number ACI-154862.

## Authors Contribution

Y.L. performed all MD simulations. Y.L.L designed and supervised the project. Y.L and Y.L.L analyzed data and wrote the paper together.

## Notes

### Competing Interest Statement

The authors have declared no competing interest.

